# CardioChrom Reconstructs the Human Cardiac Virtual Epigenome from Single-Nucleus Transcriptomes

**DOI:** 10.64898/2026.09.18.752723

**Authors:** Hongbiao Huang, Xiaodan Hui, Yingjian Li, Ahmed Abdelbaset-Ismail, Zixuan Xu, Suchen Yadav, Yi Tan

## Abstract

**Background:** Matched epigenomic profiling remains scarce in human heart failure, limiting regulatory interpretation of single-nucleus ribonucleic acid (RNA) sequencing data. We developed CardioChrom, a cardiac-specialized framework that reconstructs chromatin accessibility, histone H3 lysine 27 acetylation (H3K27ac), and histone H3 lysine 27 trimethylation (H3K27me3) from single-nucleus RNA profiles.

**Methods:** CardioChrom was developed using same-nucleus human heart datasets jointly measuring RNA with chromatin accessibility by assay for transposase-accessible chromatin (ATAC), H3K27ac, or H3K27me3, with donor-separated development and leave-one-cell-type-out testing. Frozen models were evaluated in 10-donor and 23-sample external human heart cohorts. Histone reconstruction was benchmarked against measured profiles and a Transformer. The models were then applied without refitting to an RNA-only cohort of 43 donors with heart failure with preserved ejection fraction (HFpEF).

**Results:** Across unseen cardiac cell types, CardioChrom increased area under the precision– recall curve by 0.0109 relative to within-cell-type RNA shuffling in 11 of 12 cell types and increased area under the receiver operating characteristic curve by 0.0179 in all 12.

CardioChrom exceeded controls in both external cohorts and matched or exceeded the Transformer across 6 prespecified histone metrics. In HFpEF, 3,581 donor-level pathway–layer associations met global false discovery rate, sex-adjustment, and leave-one-donor-out robustness criteria. Endothelial cells showed prominent reconstructed histone-associated changes involving coagulation, inflammation, hypoxia, and profibrotic signaling. Von Willebrand factor expression and its reconstructed regulatory states were consistently reduced, and all 6 linked candidate cis-regulatory elements were classified as repressed. Motif-supported TF–cCRE–gene hypotheses were generated in endothelial cells and fibroblasts.

**Conclusions:** CardioChrom extends RNA-only human heart failure cohorts toward cell-type-resolved virtual epigenomic and regulatory analysis while distinguishing predictions from direct measurements.

**Clinical Perspective:** *What Is New?:*

- CardioChrom is, to our knowledge, the first cardiac-specialized virtual epigenome framework to integrate same-nucleus human heart RNA–ATAC and RNA–histone reference data to reconstruct chromatin accessibility, H3K27ac, and H3K27me3 states from single-nucleus RNA profiles.
- The framework was tested beyond its training setting, including unseen donors, leave-one-cell-type-out evaluation, and independent human heart cohorts, and consistently exceeded simple RNA-shuffled or peak-frequency baselines while remaining competitive with or superior to Transformer-based comparators.

*What Are the Clinical Implications?:*

- Many heart-failure cohorts contain transcriptomic data without matched epigenomic measurements. CardioChrom provides a way to extend these RNA-only datasets toward cell-type-resolved regulatory-state analysis without requiring additional tissue or multimodal assays.
- In an independent RNA-only HFpEF cohort, CardioChrom identified prominent endothelial regulatory remodeling involving Coagulation, IL6/JAK/STAT3, TNFα/NFκB, Hypoxia, and TGF-β/EMT and generated TF–cCRE–gene hypotheses that can be prioritized for experimental validation.

## Background

Heart failure involves coordinated molecular remodeling across multiple cardiac cell types, yet most human single-cell studies remain limited to transcriptomic measurements. Chromatin accessibility and histone modifications provide complementary information about the regulatory programs underlying cardiac remodeling, but such multimodal datasets remain comparatively scarce because of limited tissue availability and additional experimental complexity (1–3).

Computational cross-modal reconstruction offers a potential solution by learning relationships from paired multiomic data and transferring them to larger RNA-only datasets. Existing approaches, including BABEL, MultiVI, scButterfly, Cisformer, SEE, and Corgi/Corgi+, have demonstrated the feasibility of predicting chromatin accessibility or other regulatory readouts from transcriptional state (4–10), supporting the emerging concept of RNA-anchored virtual regulatory-state reconstruction.

As these methods become increasingly capable, an important question is no longer simply whether one modality can be predicted from another, but whether the learned relationship remains informative across biological contexts. Models intended for scarce human tissues must generalize beyond the cells used for training, including to new individuals, cell identities, disease states, and independent studies. This is especially important for regulatory genomics, where strong locus specific and cell type associated structure may coexist with more specific information carried by the matched transcriptional state. Recent evaluations of single cell foundation models have shown that increased architectural complexity does not necessarily translate into stronger zero shot generalization, and that well specified simpler models can equal or outperform much larger models in some settings (11, 12). These findings make rigorous benchmarking and realistic transfer evaluation central components of virtual cell development.

The human heart provides a particularly informative setting for this problem because multiomic cardiac datasets remain scarce, whereas transcriptomic datasets are increasingly available. In addition, technologies such as Paired Tag and Droplet Paired Tag now enable transcriptomes and histone modifications to be measured jointly in individual nuclei (13, 14). This creates the opportunity to extend RNA based reconstruction beyond accessibility and ask whether transcriptional state can also recover complementary active and repressive chromatin landscapes represented by H3K27ac and H3K27me3.

Here, we establish an RNA anchored virtual epigenome framework for the human heart that reconstructs chromatin accessibility together with active H3K27ac and repressive H3K27me3 states from single nucleus transcriptomes. We evaluate reconstruction using donor aware development, validation, and test partitions together with leave one cell type out analysis, thereby testing generalization to both unseen individuals and unseen cardiac cell types. We benchmark compact latent state translation against nonlinear Transformer based models and use locus level priors and RNA shuffling to quantify information contributed by the matched transcriptional state. We further assess transfer across independent human heart datasets and evaluate histone reconstruction against directly measured RNA and histone pairs. Finally, the resulting virtual epigenome is applied to an independent RNA only heart failure cohort to derive cell type resolved regulatory hypotheses. By linking stringent cross modality evaluation with deployment in transcriptome only data, this framework provides a strategy for expanding the regulatory information recoverable from the growing collection of human cardiac datasets that lack matched epigenomic measurements.

## Materials and Methods

### Data Availability and Ethical Considerations

The Foundation for the National Institutes of Health (FNIH) and CAREHF multimodal human heart datasets were obtained from the study by Xie et al (3). The external GSE270788 multiome dataset was generated by Amrute et al(15), and the independent heart failure with preserved ejection fraction (HFpEF) single-nucleus RNA sequencing dataset is available through the Single Cell Portal under accession SCP3342(16). CardioChrom source code, installation instructions, and a synthetic example are available on GitHub (https://github.com/YMSWhuang/CardioChrom). The frozen model bundle is archived on Zenodo (https://doi.org/10.5281/zenodo.22239579), and the software is released under the MIT License.

### Study Design and Datasets

CardioChrom was developed as an RNA-anchored framework for reconstructing cardiac chromatin accessibility and histone states from single-nucleus transcriptomes. Model development and internal evaluation used the FNIH human heart multiomic resource, which included 329,255 paired RNA–ATAC nuclei from 30 donors and separate same-nucleus RNA–acetylation of lysine 27 on histone H3 (H3K27ac) and RNA– trimethylation of lysine 27 on histone H3 (H3K27me3) datasets containing 67,453 and 62,127 nuclei, respectively (3, 14). Donors were prespecified as 21 training, 3 validation, and 6 test donors, and 12 leave-one-cell-type-out models were used to assess generalization to unseen cardiac cell identities. CAREHF, comprising 243,008 paired RNA–ATAC nuclei from 10 donors, was used as an external transportability benchmark. GSE270788 provided an independent external RNA–ATAC confirmation dataset, with 31,477 nuclei from 23 paired samples entering the primary routed analysis (15). The frozen framework was subsequently applied to the RNA-only SCP3342 HFpEF cohort, which included 48,866 nuclei from 43 donors, including 19 patients with HFpEF and 24 nonfailing controls (16) (Figure 1). CAREHF was not used for model fitting; because CAREHF results had been inspected before the configuration was formally finalized, it was treated as a transportability benchmark rather than an untouched confirmatory cohort.

**Figure 1.**
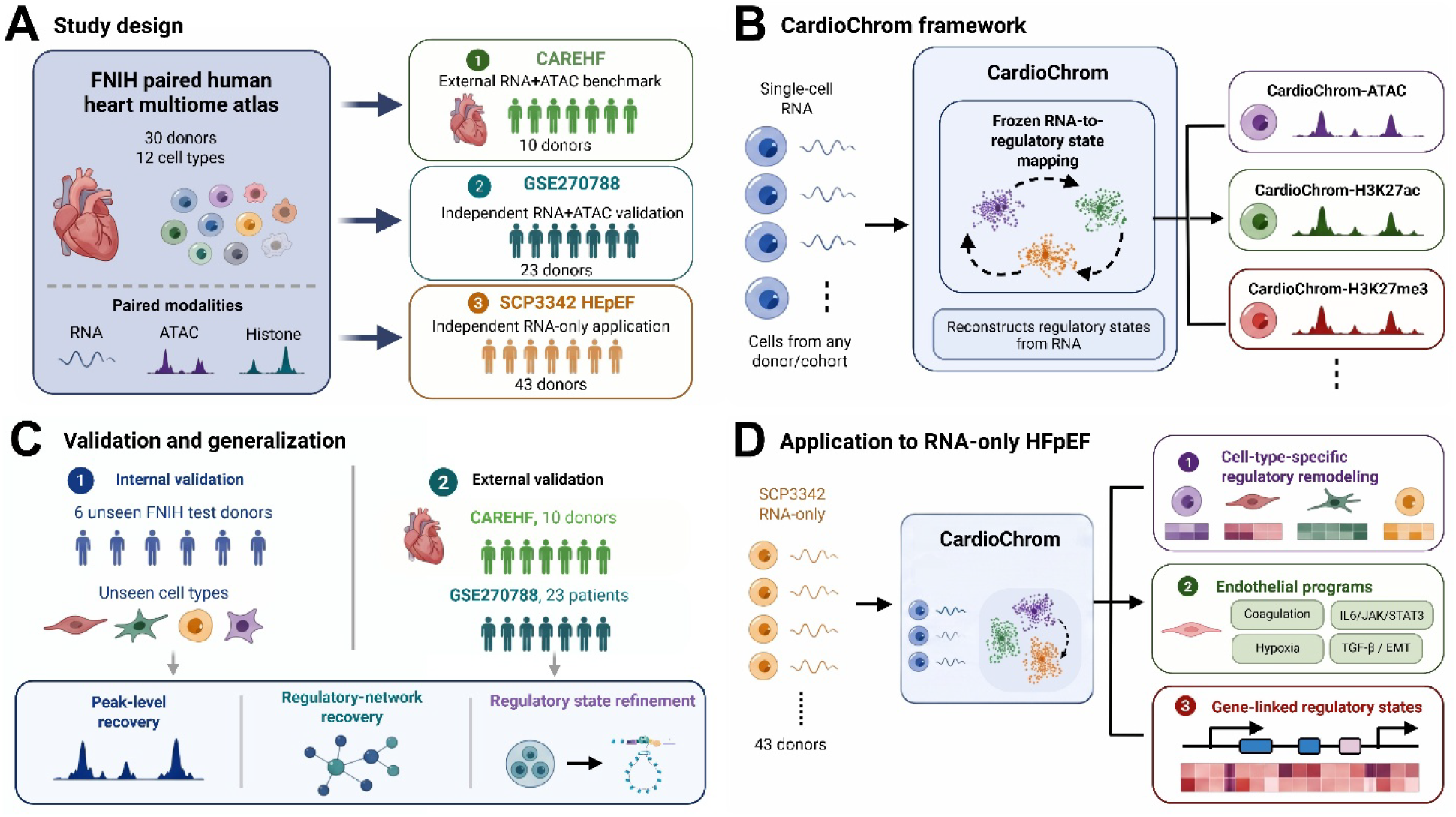
Study design A, Overview of the study cohorts and multimodal data resources. The Foundation for the National Institutes of Health (FNIH) paired human heart multiome atlas, comprising 30 donors and 12 cardiac cell types with matched RNA, ATAC, H3K27ac, and H3K27me3 measurements, was used to develop the cross-modal reconstruction framework. CAREHF was used as an external transportability benchmark, and GSE270788 as an independent confirmation cohort, whereas SCP3342 heart failure with preserved ejection fraction (HFpEF) was used as an independent RNA-only application cohort. **B,** Schematic of the CardioChrom framework. Single-nucleus RNA profiles are projected through a frozen RNA-to-regulatory-state mapping to reconstruct three virtual regulatory layers: CardioChrom-ATAC, CardioChrom-H3K27ac, and CardioChrom-H3K27me3. **C,** Internal and external validation strategy. Internal evaluation assessed generalization across six unseen FNIH test donors and held-out cell types. External validation in CAREHF and GSE270788 examined peak-level recovery, recovery of measured regulatory-network relationships, and regulatory-state refinement across independent cohorts. **D,** Application of frozen CardioChrom to the independent SCP3342 RNA-only HFpEF cohort. Reconstructed regulatory states were used to characterize cell-type-specific remodeling, define endothelial disease-associated programs including Coagulation, IL6/JAK/STAT3, Hypoxia, and TGF-β/EMT, and resolve gene-linked regulatory states at the level of individual regulatory programs.

### CardioChrom Reconstruction and Evaluation

CardioChrom was adapted from the latent-state nearest-neighbor strategy developed in the NeurIPS 2021 Multimodal Single-Cell Data Integration Challenge (17). RNA and target regulatory modalities were represented in training-derived rank-50 latent spaces. For each query nucleus, the 25 nearest training nuclei were identified from the RNA latent representation, and their target-modality latent states were averaged to reconstruct ATAC, H3K27ac, or H3K27me3 profiles. Donor identity, disease status, sex, and cohort labels were not supplied to the predictor. All feature spaces, transformations, neighborhood rules, and reconstruction procedures were fixed before final evaluation. RNA-to-ATAC reconstruction was evaluated by area under the precision–recall curve (AUPRC) and area under the receiver-operating-characteristic curve (AUROC), using a training-derived peak prior, within-cell-type RNA-shuffled control, and frozen Transformer comparator. Histone reconstruction was evaluated against directly measured same-nucleus histone profiles, with H3K27ac assessed primarily at 15-kb resolution and H3K27me3 additionally assessed using broader 75-and 225-kb representations. RNA-shuffled predictions, training-set mean profiles, and a Transformer-based DualHistFormer model served as histone comparators.

### Regulatory Network, Pathway, and HFpEF Analyses

Regulatory networks were inferred with SCENIC+ from measured ATAC, CardioChrom-reconstructed ATAC, and RNA-shuffled ATAC using matched RNA profiles (18). Recovery of measured region-to-gene and transcription factor (TF)–region–gene relationships was evaluated using matched top-ranked network sets. Source-derived activity-by-contact (ABC) links were used to connect regulatory features to genes (19). Gene-level regulatory effects were analyzed using preranked gene-set enrichment analysis with Hallmark and Reactome gene sets (20–22). For SCP3342, frozen CardioChrom models were applied without refitting or recalibration.

Donor-level disease effects were calculated across measured RNA, reconstructed ATAC and histone layers, and derived histone and regulatory composite scores. RNA-only transcription factor activity was inferred with pySCENIC, and CardioChrom transcription factors were prioritized from motif-supported direct TF–gene relationships and reconstructed disease effects; motif-resolved TF–cCRE–gene networks incorporated JASPAR motif support (23, 24).

### Statistical Analysis and Human Subjects Oversight

Donors or external biological samples, rather than individual nuclei or genomic features, were treated as the units of statistical inference. Paired external-cohort comparisons used 2-sided Wilcoxon signed-rank tests. HFpEF-associated gene and pathway effects were summarized using Hedges’ *g*, with Benjamini–Hochberg correction for multiple testing (25). Robust pathway associations required global false discovery rate (FDR) <0.05 in both primary and sex-adjusted analyses and concordant effect direction in at least 90% of leave-one-donor-out analyses. The University of Arizona Institutional Review Board determined that this secondary analysis of deidentified data did not constitute human subjects research (STUDY00008718). Detailed preprocessing, sampling procedures, model specifications, evaluation metrics, sensitivity analyses, statistical models, software versions, and random seeds are provided in the Supplemental Methods.

## Results

### CardioChrom reconstructs chromatin accessibility from RNA across unseen cardiac cell types and donors

We first evaluated whether matched transcriptional state contained reproducible information for reconstructing chromatin accessibility beyond locus-level accessibility priors and cell-type-associated structure. CardioChrom was evaluated across 12 leave-one-cell-type-out folds in the FNIH cardiac multiome cohort (Figure 2A). Across held-out cell types, predictions generated from matched RNA profiles generally outperformed the within-cell-type RNA-shuffled control. Relative to RNA shuffling, CardioChrom increased AUPRC by a mean of 0.0109, with positive gains in 11 of 12 held-out cell types, and increased AUROC by 0.0179, with positive gains in all 12 cell types (Figure 2A and 2B).

**Figure 2.**
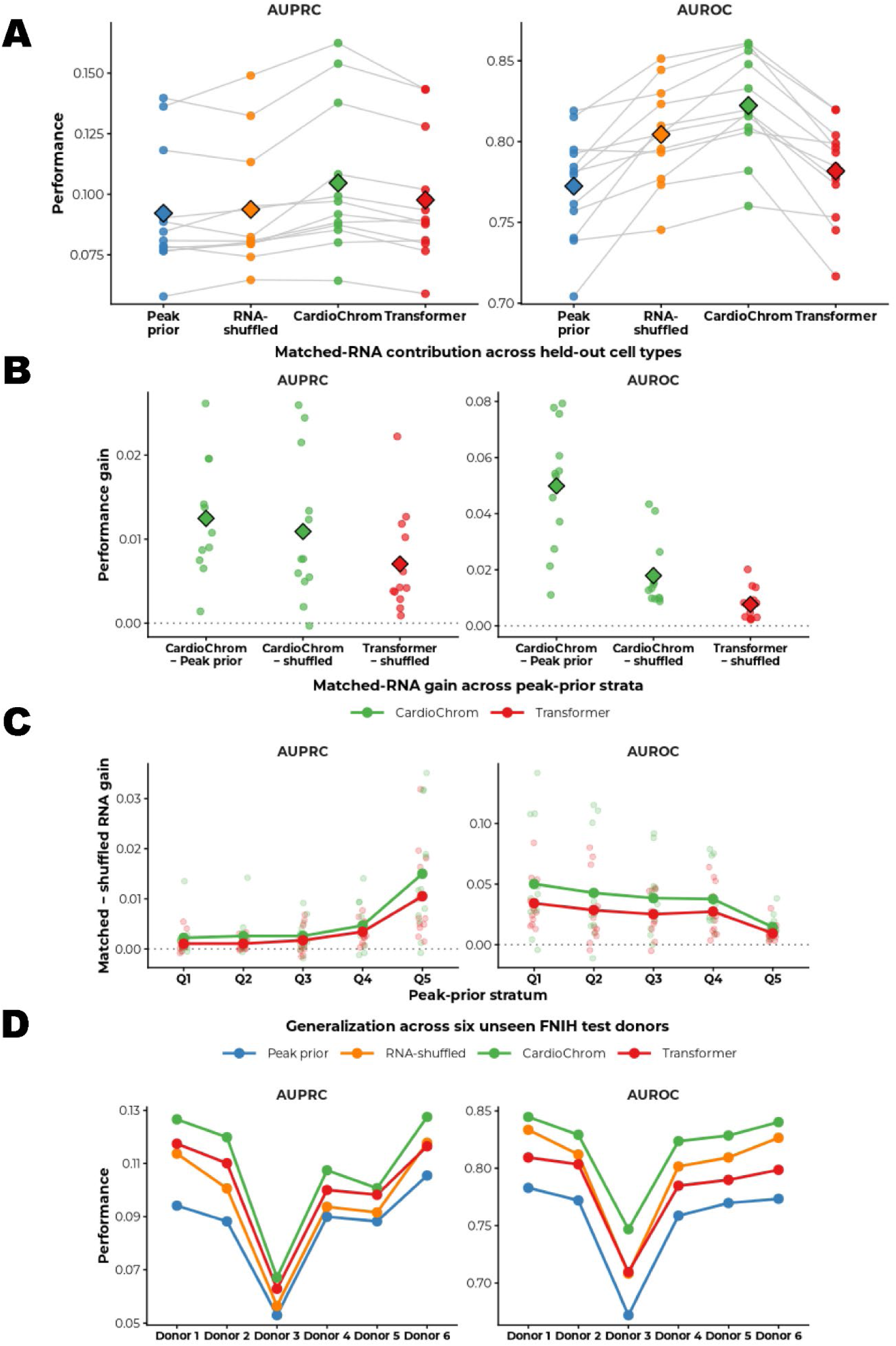
Internal validation of RNA-to-chromatin accessibility prediction in the Foundation for the National Institutes of Health (FNIH) cardiac multiome cohort A, Performance of the frozen RNA-to-ATAC models across 12 leave-one-cell-type-out folds, evaluated by area under the precision–recall curve (AUPRC) and area under the receiver operating characteristic curve (AUROC). Each colored point represents one held-out cell type, gray lines connect results from the same held-out cell type across model configurations, and diamonds indicate the mean across the 12 cell types. Peak prior represents the accessibility prior without RNA information; RNA-shuffled serves as a negative control in which the correspondence between RNA profiles and ATAC targets was disrupted; CardioChrom denotes prediction using the matched RNA input; Transformer denotes the frozen Transformer-based comparator. **B,** Contribution of matched RNA information across the 12 held-out cell types. Performance gains are shown for CardioChrom relative to the peak prior, CardioChrom relative to its RNA-shuffled control, and Transformer relative to its corresponding shuffled-RNA control. Each point represents one held-out cell type and diamonds indicate the mean gain. Positive values indicate improved predictive performance from the corresponding RNA-informed model. **C,** Matched-RNA gain stratified by peak-prior quintile (Q1–Q5). Points show cell-type-specific gains in AUPRC or AUROC relative to the corresponding shuffled-RNA control, and solid lines show the mean across the 12 held-out cell types for CardioChrom and Transformer models. Q1 represents peaks with the lowest prior accessibility and Q5 those with the highest prior accessibility. **D,** Generalization across six completely unseen FNIH test donors. For each donor, AUPRC and AUROC were macro-averaged across the 12 cardiac cell types. Each donor–model combination therefore contributes one value per metric. The six held-out donors are displayed as Donor 1– Donor 6. Across panels, blue denotes Peak prior, orange RNA-shuffled, green CardioChrom, and red Transformer.

The contribution of matched RNA was retained across strata of baseline peak accessibility (Figure 2C), indicating that reconstruction was not explained solely by learning frequently accessible genomic regions. A frozen Transformer comparator did not provide a consistent improvement over the compact latent-state transfer model.

We further assessed generalization to six FNIH donors completely excluded from model development. CardioChrom maintained stable AUPRC and AUROC across the six held-out donors after macro-averaging across cardiac cell types (Figure 2D), supporting reproducible accessibility reconstruction across unseen cell types and individuals.

### CardioChrom generalizes across independent human heart cohorts and recovers regulatory-network structure

We next tested whether the frozen RNA-to-ATAC mapping could transfer to independent human heart datasets without cohort-specific retraining or recalibration. The FNIH-derived CardioChrom model was first applied to CAREHF, comprising 243,008 paired RNA–ATAC nuclei from 10 donors (3) (Figure 3A). Across 12 cardiac cell types, CardioChrom showed higher overall AUPRC and AUROC than the peak prior and RNA-shuffled control and also outperformed the frozen Transformer comparator on average (Figure 3B).

**Figure 3.**
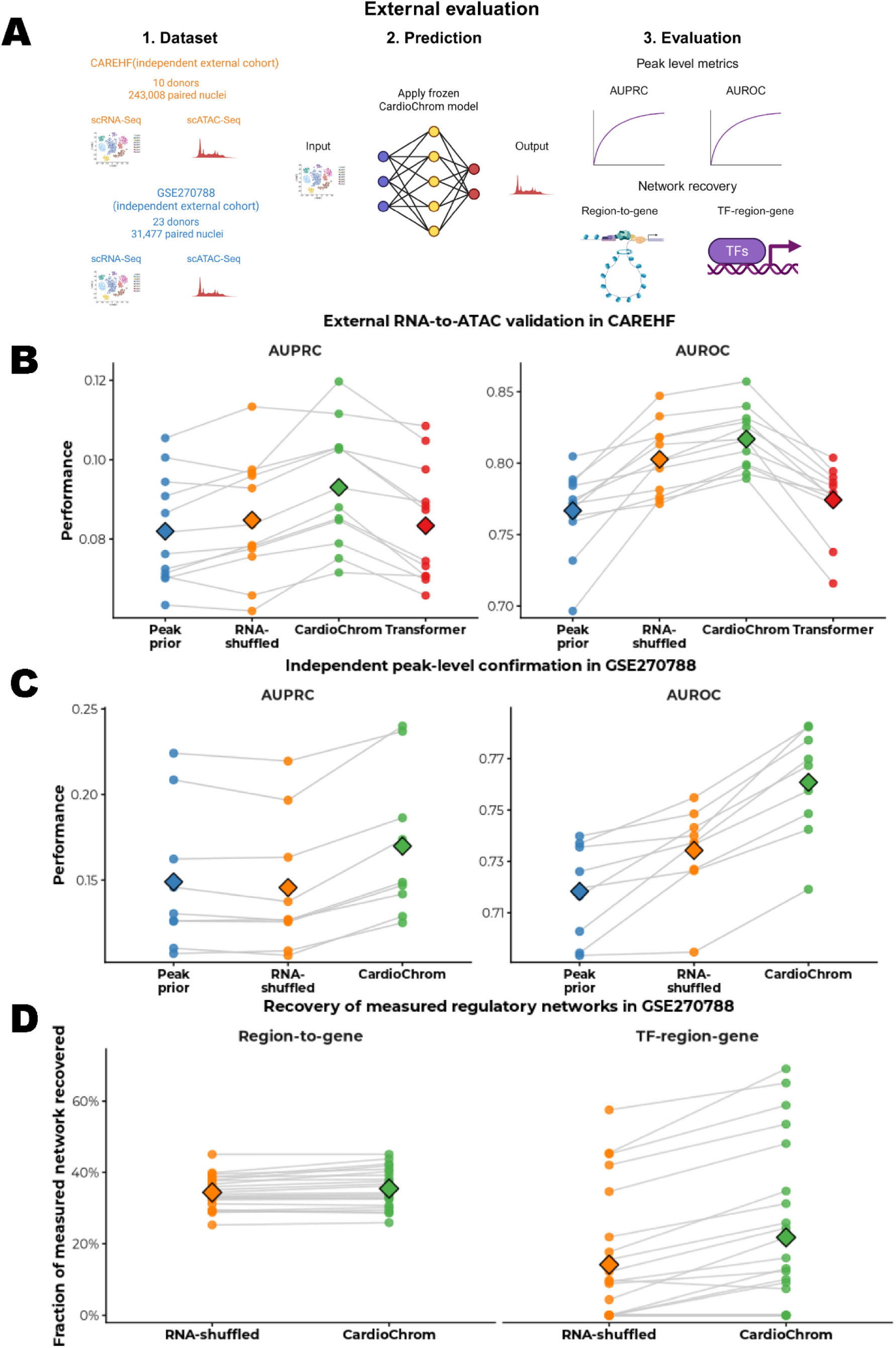
External validation and regulatory-network recovery of RNA-to-ATAC reconstruction A, External RNA-to-ATAC evaluation framework. The Foundation for the National Institutes of Health (FNIH)-derived model was applied without retraining or external model fitting to CAREHF (10 donors; 243,008 paired RNA–ATAC nuclei) and GSE270788 (23 samples; 31,477 primary-routed paired nuclei). Peak-level performance was assessed by area under the precision– recall curve (AUPRC) and area under the receiver operating characteristic curve (AUROC), and GSE270788 was additionally used for regulatory-network recovery. **B,** External RNA-to-ATAC evaluation in CAREHF across 12 cardiac cell types. Peak prior, RNA-shuffled, CardioChrom, and Transformer performance is shown for AUPRC and AUROC. Points represent cell types, gray lines connect matched cell types, and diamonds indicate the mean. **C,** Independent peak-level confirmation in GSE270788 across nine routed cell types. CardioChrom was compared with the peak prior and RNA-shuffled control using measured ATAC as the reference. Points represent cell types, gray lines connect matched results, and diamonds indicate the mean. **D,** Regulatory-network recovery in 23 GSE270788 samples. Region-to-gene and transcription factor (TF)–region–gene relationships reconstructed from CardioChrom and RNA-shuffled virtual ATAC were compared with networks derived from measured multiome data. Points represent samples, gray lines connect paired results, and diamonds indicate the mean. Recovery was evaluated using the top 5% of ranked regulatory features.

A second external evaluation was performed in GSE270788 (15), in which 31,477 nuclei from nine prespecified cardiac cell-type routes entered the primary analysis. Using measured ATAC as the reference, CardioChrom achieved higher AUPRC and AUROC than both controls in all nine routed cell types (Figure 3C), supporting transfer across distinct donor populations, disease contexts, and experimental datasets.

We then assessed whether reconstructed accessibility preserved downstream regulatory information. Region-to-gene relationships and transcription factor–region–gene triplets were reconstructed from CardioChrom and RNA-shuffled virtual ATAC profiles across the 23 GSE270788 samples and compared with networks inferred from measured multiome data.

CardioChrom recovered a greater fraction of top-ranked measured relationships for both network levels, with a more pronounced separation for TF–region–gene triplets (Figure 3D).

### RNA-anchored reconstruction captures active and repressive histone states

We next asked whether CardioChrom could extend RNA-anchored reconstruction beyond chromatin accessibility to active and repressive histone states. Separate H3K27ac and H3K27me3 models were developed using same-nucleus RNA–histone measurements from the FNIH resource (3), without pseudo-pairing across histone marks. In the held-out measured-histone benchmark, both CardioChrom and the within-cell-type RNA-shuffled control showed greater genome-wide donor-pseudobulk agreement with measured histone profiles than the training-set mean, although the two RNA-based reconstructions were similar at this whole-genome level (Figure 4A). H3K27ac was evaluated at 15-kb resolution, whereas H3K27me3 used a prespecified broad-domain metric integrating 75-and 225-kb scales.

**Figure 4.**
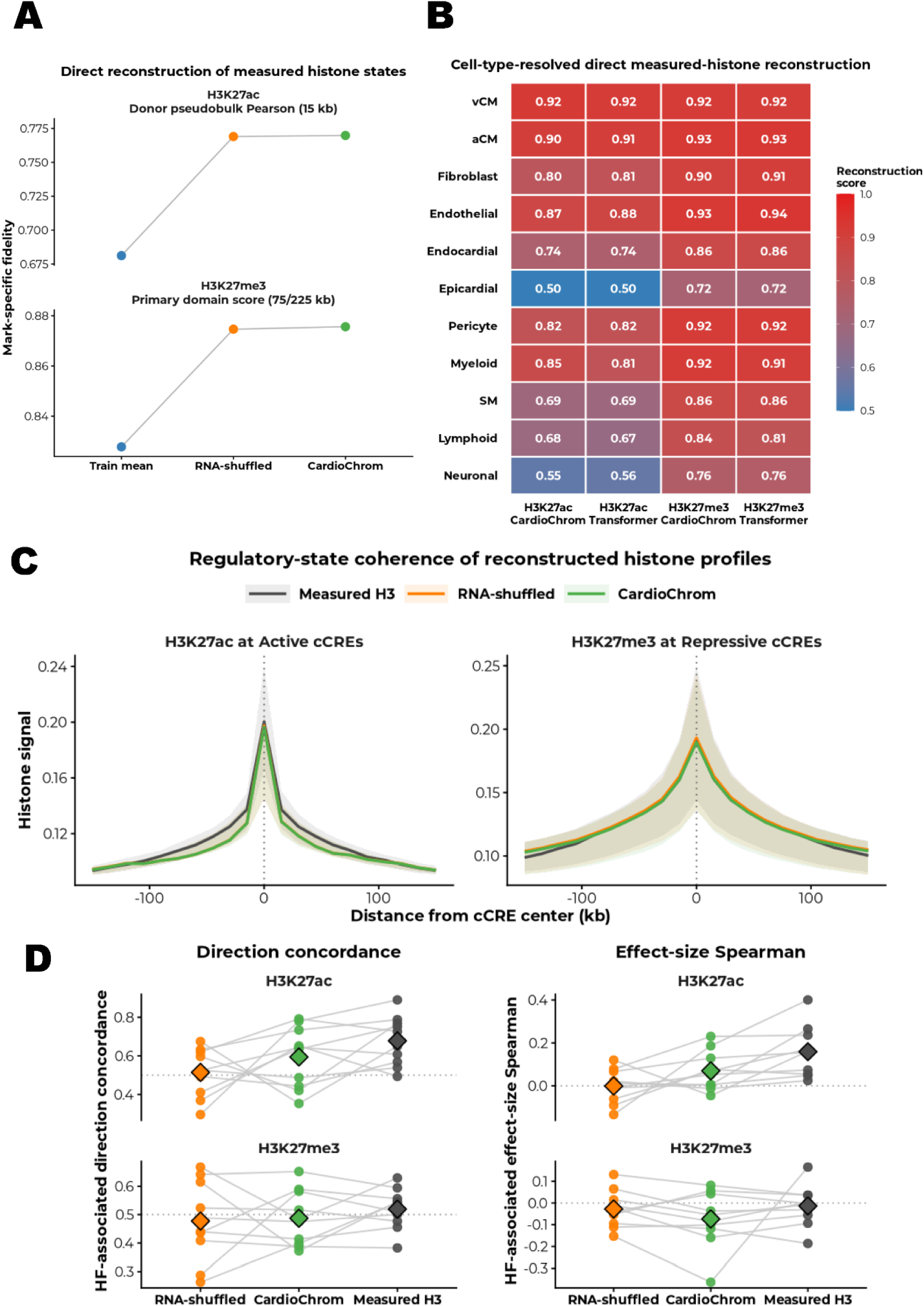
CardioChrom reconstructs active and repressive histone states across cardiac cell types and preserves disease-associated chromatin remodeling A, Direct reconstruction of measured histone states. H3K27ac was evaluated by donor-pseudobulk Pearson correlation at 15-kb resolution, whereas H3K27me3 was evaluated using a broad-domain score integrating 75-and 225-kb scales. Train mean, RNA-shuffled, and CardioChrom predictions are shown. **B,** Cell-type-resolved reconstruction in held-out cardiac cell types. Heatmaps compare CardioChrom and Transformer performance for H3K27ac and H3K27me3. H3K27ac values represent donor-pseudobulk Pearson correlations at 15-kb resolution; H3K27me3 values represent the broad-domain reconstruction score. **C,** Regulatory-state coherence around cardiac cis-regulatory elements. H3K27ac profiles were centered on active cCREs and H3K27me3 profiles on repressive cCREs. Lines show median signal across donor–cell-type units and shaded regions indicate the interquartile range. Gray, orange, and green denote measured H3, RNA-shuffled, and CardioChrom, respectively. **D,** Recovery of heart failure (HF)-associated chromatin remodeling across cardiac cell types. Disease-associated histone changes were compared with corresponding measured ATAC changes. Direction concordance is shown on the left and orientation-adjusted effect-size Spearman correlation on the right; H3K27me3 was sign-adjusted for its inverse relationship with accessibility. Points represent cell types, gray lines connect matched results, and diamonds indicate the mean. Dotted lines indicate reference values of 0.5 for direction concordance and 0 for Spearman correlation.

We next compared CardioChrom with DualHistFormer, a Transformer-based nonlinear comparator. Across six prespecified metrics spanning genome-wide agreement, recovery of high-signal regions, and heart-failure-associated differential effects, CardioChrom matched or exceeded the Transformer on all six measures (Supplemental Figure 1).

Evaluation across held-out cardiac cell types further demonstrated reconstruction in cell identities excluded from model fitting (Figure 4B). Performance was strongest in ventricular and atrial cardiomyocytes and endothelial cells, with H3K27ac Pearson correlations of 0.92, 0.90, and 0.87 and corresponding H3K27me3 broad-domain scores of 0.92, 0.93, and 0.93, respectively. Fibroblast, pericyte, and myeloid compartments also showed strong recovery, whereas lymphoid, epicardial, and neuronal compartments showed lower performance.

Reconstructed profiles also retained the expected cCRE-centered spatial organization, with a sharp H3K27ac peak around active cCREs and a broader H3K27me3 distribution around repressive cCREs (Figure 4C).

Finally, we examined whether reconstructed histone states preserved heart failure-associated chromatin remodeling. Histone disease effects were compared with corresponding measured ATAC effects across held-out cell types, with H3K27me3 orientation-adjusted for its expected inverse relationship with accessibility. Recovery was strongest for H3K27ac, whereas H3K27me3 showed more modest cross-assay concordance (Figure 4D).

### Virtual histone states recover disease-associated regulatory programs across heart-failure cohorts

Having established reconstruction against measured histone profiles, we next examined whether virtual histone states could recover disease-associated regulatory programs. Thirteen representative pathways were selected using predefined criteria requiring CardioChrom to recover the measured-H3 direction while the RNA-shuffled control did not and to yield an normalized enrichment score (NES) closer to measured H3. After collapsing highly overlapping pathways, six ventricular-cardiomyocyte, four endothelial, and three pericyte programs were retained (Supplemental Figure 2A). Recovery was most prominent in ventricular cardiomyocytes, including Interleukin-1 Family Signaling, Biological Oxidations, KEAP1– NFE2L2 signaling, and Selective Autophagy; endothelial and pericyte programs included DNA repair, mitochondrial regulation, extracellular-matrix organization, metabolism, and remodeling.

We then asked whether virtual histone states could refine measured chromatin accessibility in CAREHF. Because CAREHF lacked measured histone profiles, agreement was assessed against the FNIH measured-H3 pathway reference rather than direct histone accuracy. The strongest gains occurred in cardiomyocytes (Supplemental Figure 2B). In ventricular cardiomyocytes, adding virtual H3 to measured ATAC increased Reactome pathway-pattern Spearman correlation by 0.655, reduced median absolute NES error by 1.429, increased direction agreement by 0.46, and increased pathway-recovery fraction by 0.12. Corresponding atrial-cardiomyocyte gains were 0.549, 0.416, 0.30, and 0.28. Hallmark results were more modest or mixed, supporting context-dependent rather than uniform refinement.

We next used GSE270788 to test recovery of an experimentally supported disease program from the source study (15). The original study identified IL-1β signaling from CCR2+ monocytes/macrophages to fibroblasts as a driver of the FAP/POSTN profibrotic state and validated this axis in vivo. We therefore focused on the fibroblast arm. In cardiomyopathy fibroblasts, integration of measured ATAC with virtual H3 yielded a more positive Interleukin-1 Family Signaling NES than measured RNA or ATAC alone (Figure 5A). FAP, IL1R1, and MEOX1 showed positive disease-associated RNA signals, and their reconstructed regulatory-composite scores remained positive (Figure 5B). The macrophage ligand arm was not evaluated because the external crosswalk lacked a CCR2+ macrophage-specific population.

**Figure 5.**
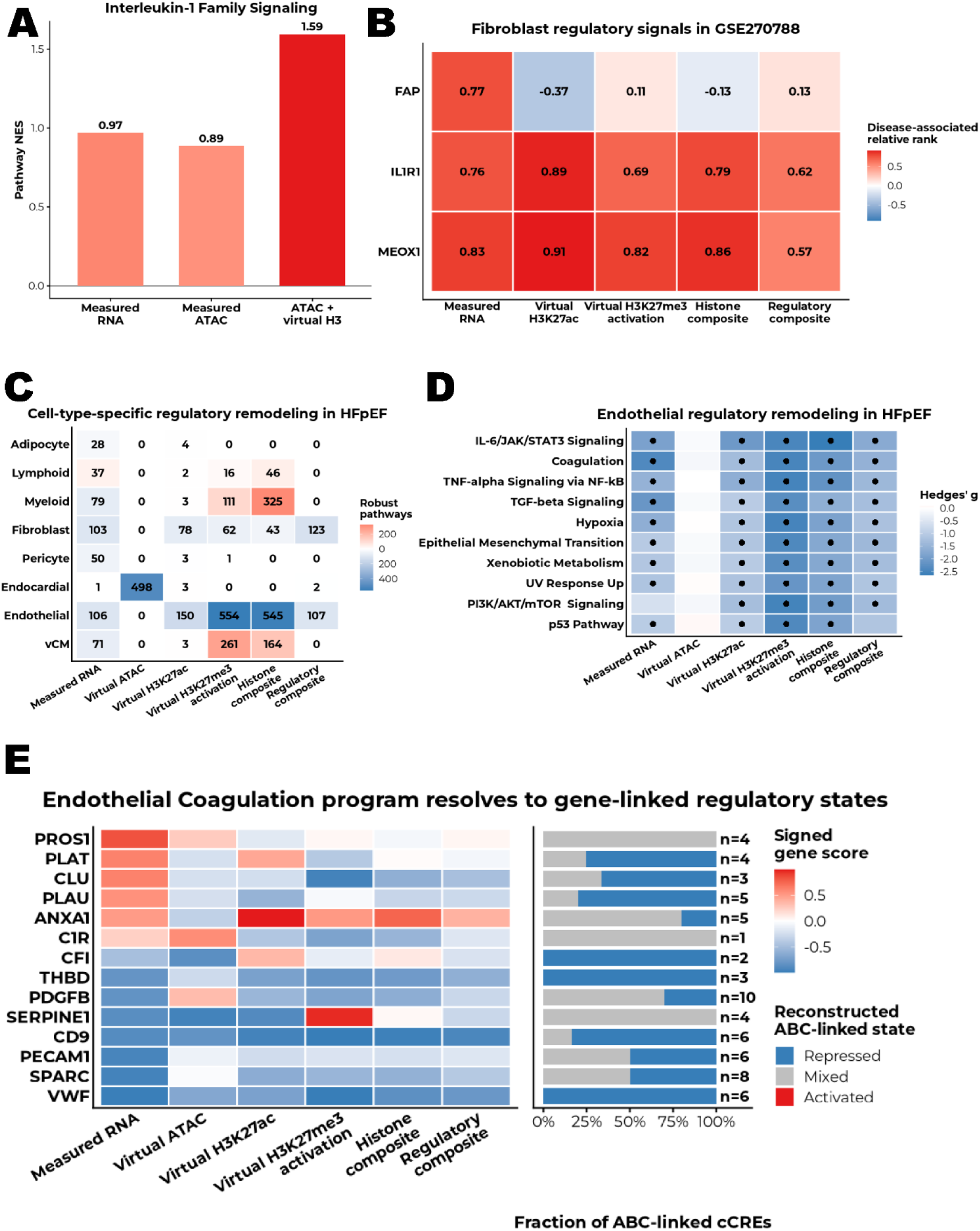
Biological validation and disease-state reconstruction with CardioChrom across heart-failure cohorts A, Recovery of Interleukin-1 Family Signaling in GSE270788 fibroblasts. Bars show pathway normalized enrichment score (NES) across measured RNA, measured ATAC, and ATAC + virtual H3. **B,** Disease-associated regulatory states of selected fibroblast genes in GSE270788. Tiles show relative ranks across measured RNA, virtual H3K27ac, virtual H3K27me3 activation, histone composite, and regulatory composite layers. **C,** Cell-type-specific regulatory remodeling in the SCP3342 heart failure with preserved ejection fraction (HFpEF) cohort. Numbers indicate the number of robust HFpEF-associated pathway effects; tile color indicates the predominant direction. **D,** Endothelial regulatory remodeling in HFpEF. The heatmap shows the top 10 Hallmark pathways across measured RNA and reconstructed regulatory layers. Colors indicate Hedges’ *g*, and black dots denote robust pathway effects. **E,** Gene-level decomposition of the endothelial Coagulation program. The left heatmap shows signed disease-associated gene scores across measured and reconstructed layers. Stacked bars show the proportions of activity-by-contact (ABC)-linked candidate cis-regulatory elements (cCREs) classified as Repressed, Mixed, or Activated; *n* indicates the number of linked cCREs.

### Frozen deployment in RNA-only HFpEF reveals prominent endothelial regulatory remodeling

We next deployed the frozen CardioChrom models in SCP3342, an independent RNA-only HFpEF cohort containing 48,866 nuclei from 43 donors, including 19 patients with HFpEF and 24 nonfailing controls (16). Without refitting or recalibration, 48,433 nuclei from 10 cardiac cell types were mapped to predefined CardioChrom reference cell types and reconstructed across virtual ATAC, H3K27ac, and H3K27me3 states; 433 lymphatic endothelial nuclei remained unmapped (Figure 5C).

Donor-level analyses across measured and reconstructed regulatory layers identified 3,581 robust pathway–layer effects meeting global FDR significance in both primary and sex-adjusted analyses and concordant direction in at least 90% of leave-one-donor-out analyses. Endothelial cells showed the most prominent histone-associated remodeling, with 554 robust effects in virtual H3K27me3 activation, 150 in virtual H3K27ac, and 545 in the histone composite layer (Figure 5C). Other compartments showed distinct patterns, including prominent accessibility-associated remodeling in endocardial cells and more selective signals in fibroblasts.

We next resolved the endothelial signal at the pathway level. The 10 highest-ranked Hallmark programs included Coagulation, IL6/JAK/STAT3 Signaling, TNFα Signaling via NFκB, Hypoxia, TGF-β Signaling, and Epithelial Mesenchymal Transition (Figure 5D). Across several programs, lower measured RNA activity was accompanied by lower predicted H3K27ac and H3K27me3 activation scores, whereas accessibility changes were often weaker, localizing a substantial component of HFpEF-associated endothelial remodeling to reconstructed histone states.

Coagulation was selected for gene-level resolution because of its relevance to endothelial hemostatic and vascular-homeostasis biology (Figure 5E). Several vascular genes showed concordant transcriptional and reconstructed regulatory suppression. VWF was particularly consistent: measured RNA and each reconstructed regulatory layer decreased in HFpEF, and all six source-defined ABC-linked cCREs were classified as repressed. This pattern agreed with the original SCP3342 study (16) and an independent high-fat diet (HFD)/L-NAME mouse HFpEF study reporting reduced cardiac Vwf expression with impaired microvascular density (26).

### CardioChrom reprioritizes cell-type-specific transcriptional regulators and reconstructs motif-supported regulatory networks

Finally, we asked whether CardioChrom could identify regulatory organization beyond RNA-only analysis. We focused on endothelial cells and fibroblasts, two compartments with prominent remodeling across multiple regulatory layers (Figure 5C), and compared CardioChrom TF prioritization with RNA-only pySCENIC in the same SCP3342 cohort.

CardioChrom identified distinct cell-type-specific regulatory repertoires (Figure 6A and 6C). The highest-ranked endothelial regulators were ETS2, SOX18, FLI1, EBF1, and ELF1, whereas fibroblast prioritization was led by NFE2L1, KLF5, FOSL2, KLF4, and FOSB.

**Figure 6.**
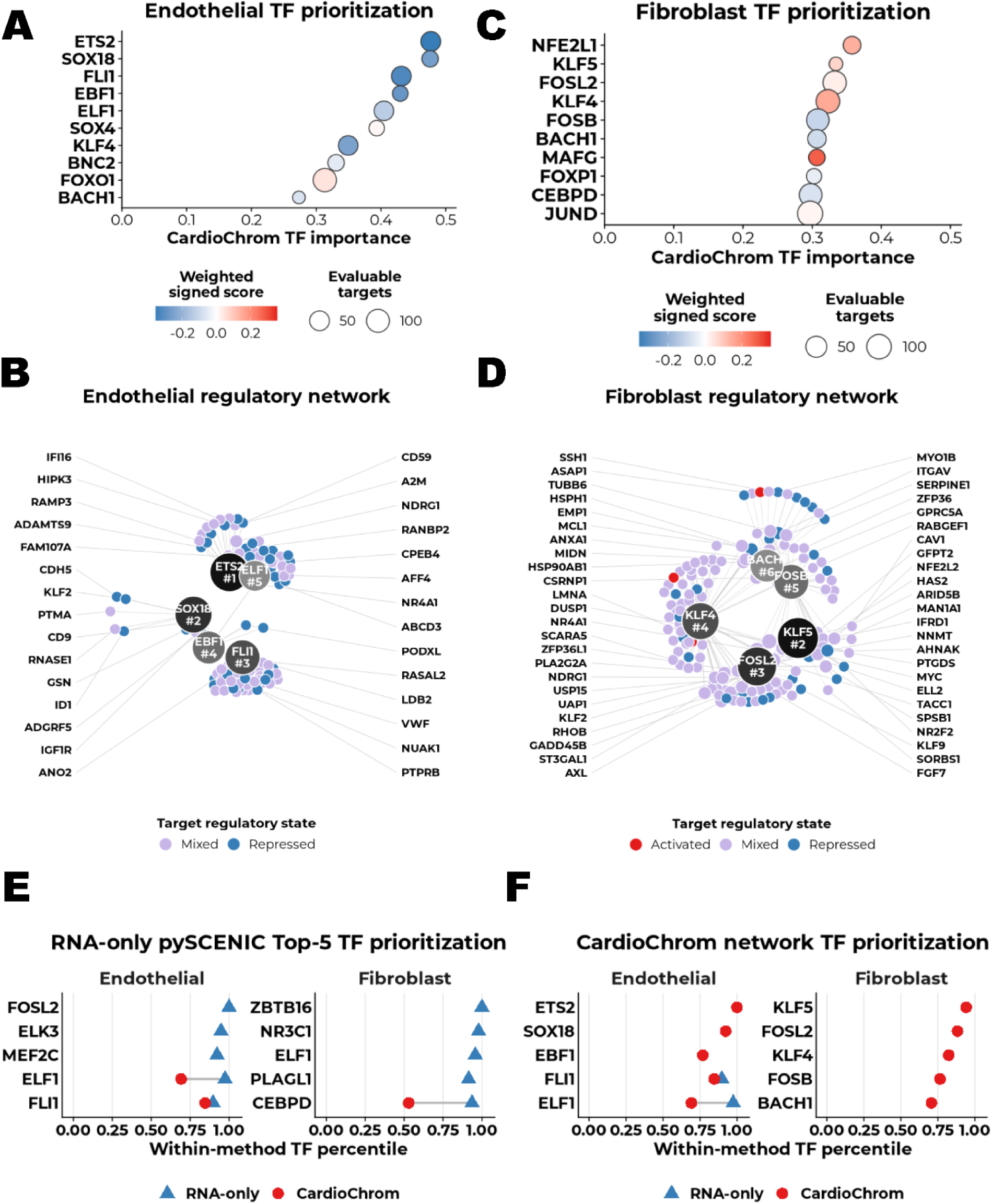
Cell-type-specific transcription factor prioritization and regulatory networks in heart failure with preserved ejection fraction (HFpEF) (A, **C)** CardioChrom-based transcription factor (TF) prioritization in endothelial cells and fibroblasts, respectively. TFs are ranked by CardioChrom importance; bubble size indicates the number of evaluable target genes and color indicates the weighted signed regulatory score. **(B, D)** Reconstructed TF–target regulatory networks in endothelial cells and fibroblasts, respectively. Large nodes indicate prioritized TFs, and target genes are colored by reconstructed HFpEF regulatory state: activated, mixed, or repressed. **(E)** Comparison of the top five RNA-only pySCENIC TFs with their CardioChrom percentile ranks. Blue triangles indicate RNA-only pySCENIC and red circles indicate CardioChrom. NR indicates not represented in the CardioChrom-eligible TF set. **(F)** Comparison of CardioChrom-prioritized TFs with their RNA-only pySCENIC percentile ranks. Symbols are as in panel E. NR indicates not represented among motif-pruned RNA-only pySCENIC regulons.

We next required complete evidence chains linking prioritized TFs to direct TF–gene relationships, cardiac ABC-linked cCREs, HFpEF-associated reconstructed regulatory states, and cCRE-specific motif support. Inferred networks were reconstructed for ETS2, SOX18, FLI1, EBF1, and ELF1 in endothelial cells and for KLF5, FOSL2, KLF4, FOSB, and BACH1 in fibroblasts (Figure 6B and 6D). NFE2L1, although ranked first in fibroblasts, was not displayed because it lacked a complete cCRE-specific motif-supported evidence chain.

RNA-only pySCENIC and CardioChrom showed limited overlap and substantial regulator reprioritization (Figure 6E and 6F). Several highly ranked TFs were weakly ranked or not represented by the other approach, indicating that CardioChrom highlighted candidate regulators and motif-supported target structures not prominent in RNA-only regulon activity.

## Discussion

In this study, we developed CardioChrom as a cardiac-specialized, RNA-anchored virtual epigenome framework for reconstructing chromatin accessibility together with active H3K27ac and repressive H3K27me3 states. Rather than seeking greater architectural complexity, the framework was designed around a practical problem in human cardiovascular research: transcriptomic datasets are becoming increasingly available, whereas matched single-cell or single-nucleus measurements of accessibility, histone modifications, and other regulatory modalities remain comparatively scarce. CardioChrom therefore uses information learned from deeply profiled human hearts to extend the regulatory interpretation of RNA-only cardiac datasets. Its principal contribution lies in the combination of organ-specific cross-modal learning, stringent transfer evaluation, and downstream disease-state interpretation rather than in the introduction of a new large neural architecture.

Cross-modality translation is an increasingly active area of single-cell genomics. BABEL, MultiVI, scButterfly, and Cisformer established that relationships between RNA and chromatin accessibility can be learned computationally, while more recent context-aware approaches such as Corgi/Corgi+ have extended prediction toward broader epigenomic readouts (4–7, 10).

However, generalization to biological contexts that are absent or sparsely represented during training remains a central challenge. Recent benchmarking studies have shown that larger foundation models do not consistently outperform simpler approaches in zero-shot settings and that performance can depend strongly on the composition of the training data (11, 12, 27). These observations motivate a complementary strategy to broadly pretrained models: a domain-focused model trained and tested in the biological system in which it is intended to be used. For the heart, this is particularly relevant because access to diseased human myocardium remains limited and deeply matched multiomic datasets are still uncommon (28).

An important feature of CardioChrom is that its histone mappings were learned from same-nucleus RNA–H3K27ac and RNA–H3K27me3 measurements generated in human cardiac tissue. Joint profiling reduces the ambiguity introduced when transcriptional and epigenomic measurements must be computationally matched across different cells, samples, or studies, and provides a direct reference for learning cell-state relationships. The FNIH resource therefore offered an unusual opportunity to evaluate whether RNA-associated regulatory information extends beyond accessibility to active and repressive histone organization. At the same time, our results emphasize that increased model complexity is not itself sufficient for better biological transfer: the compact latent-state framework remained competitive with or superior to the Transformer comparators across the prespecified evaluations. This finding is consistent with emerging evidence that rigorous data partitioning and biologically appropriate evaluation may matter as much as model scale for out-of-distribution single-cell prediction.

The purpose of virtual epigenomic reconstruction is not to create molecular information that is independent of the transcriptome. Rather, CardioChrom reorganizes RNA-derived cell state through cross-modal relationships learned from directly paired cardiac data. This distinction is important for interpreting both the histone reconstructions and downstream regulatory networks. In independent cohorts, virtual histone states selectively refined pathway-level interpretation rather than uniformly improving every cell type or pathway collection. Likewise, the divergence between RNA-only pySCENIC and CardioChrom TF prioritization suggests that reconstructed regulatory states can expose regulatory hypotheses that are not prominent when transcriptional regulon activity is considered alone. By connecting candidate TFs to disease-associated target states, cardiac cCRE–gene relationships, and motif support, CardioChrom converts an RNA-only disease signal into experimentally testable regulatory hypotheses rather than replacing direct epigenomic or perturbational measurements.

The HFpEF application illustrates the potential value of this strategy in a setting where only single-nucleus transcriptomes were available. Frozen deployment localized prominent reconstructed histone-associated remodeling to endothelial cells and subsequently resolved this signal from the cell-type landscape to vascular and inflammatory pathways and then to gene-and cCRE-linked regulatory states. The endothelial Coagulation program was particularly informative because its measured transcriptional direction agreed with the source HFpEF study, including reduced VWF expression, while the reconstructed layers suggested a broader alteration of endothelial hemostatic and vascular-homeostasis regulation (16). This should not be interpreted as systemic hypocoagulability; rather, it highlights the distinction between myocardial endothelial transcriptional state and circulating hemostatic phenotypes. More broadly, the endothelial findings are consistent with extensive evidence implicating coronary microvascular inflammation, rarefaction, oxidative stress, and endothelial dysfunction in HFpEF (29–34). CardioChrom extends this literature by providing candidate regulatory architectures that can be prioritized for subsequent experimental study.

Several limitations define the appropriate scope of these conclusions. CardioChrom is a virtual epigenome framework rather than a complete dynamic virtual cell, and reconstructed chromatin states remain predictions conditioned on RNA and the cardiac training reference. The SCP3342 cohort lacks directly measured ATAC or histone profiles, preventing direct epigenomic validation in HFpEF, and some external cell populations could not be mapped to a prespecified cardiac reference route. Generalization to very rare, previously unseen, or substantially altered cell states is therefore likely to remain challenging, as it is for other cross-modal and foundation-model approaches. In addition, the regulatory networks are computationally inferred and require perturbational, biochemical, or spatial validation. Future work should expand matched cardiac multiomics across diseases, ancestries, developmental states, and vascular compartments and should prospectively test prioritized TF–cCRE–gene relationships. In this sense, CardioChrom represents one step toward an organ-specialized cardiovascular virtual cell in which experimentally accessible transcriptomic measurements can be connected to increasingly rich regulatory-state models and used to guide targeted experiments(35).

## Author contributions

H.H. and Y.T. conceived and designed the study. H.H. wrote the manuscript. Y.L., X.H., A.I., Z.X., S.Y. and Y.T. edited the manuscript. Y.T. supervised this study. All authors have approved the final version of the manuscript.

## Data Access and Responsibility

H.H. had full access to all the data in the study and takes responsibility for the integrity of the data and the accuracy of the data analysis.

