## Supplemental Methods for "CardioChrom Reconstructs the Human Cardiac Virtual Epigenome from Single-Nucleus Transcriptomes"

### Materials and Methods

#### Study Design and Analytical Framework

CardioChrom was developed as an RNA-anchored framework for reconstructing cell-type-resolved chromatin accessibility and histone-modification profiles from single-nucleus RNA sequencing data. The study comprised four analytically distinct stages: model development and internal evaluation in the Foundation for the National Institutes of Health heart-failure resource; external assessment in the CAREHF cohort; untouched external confirmation using GSE270788; and frozen-model application to the RNA-only SCP3342 heart failure with preserved ejection fraction cohort. External datasets were not used to refit feature spaces, latent transformations, nearest-neighbor parameters, prediction heads, or downstream significance thresholds.

The FNIH RNA–ATAC dataset contained 329,255 paired nuclei from 30 donors, including 10 nonfailing controls, 10 patients with ischemic cardiomyopathy, and 10 patients with nonischemic cardiomyopathy (1). Twelve source-defined cardiac cell types were analyzed: ventricular cardiomyocytes, atrial cardiomyocytes, adipocytes, fibroblasts, endothelial cells, endocardial cells, epicardial cells, pericytes, myeloid cells, smooth muscle cells, lymphoid cells, and neuronal cells. Donors were partitioned into 21 training, 3 validation, and 6 test donors with disease groups balanced across partitions. All data from the test donors remained unavailable during model fitting and model selection.

Same-nucleus RNA–histone datasets from the same resource comprised 67,453 RNA–H3K27ac nuclei and 62,127 RNA–H3K27me3 nuclei generated by droplet-based Paired-Tag(1, 2). H3K27ac and H3K27me3 models were trained and evaluated separately. No pseudo-pairing or cross-mark matching of nuclei was performed.

CAREHF contained 243,008 paired RNA–ATAC nuclei from 10 donors, comprising 5 controls and 5 patients with ischemic heart failure(1). CAREHF did not contribute to model fitting. The final CardioChrom configuration was selected on the basis of the internal FNIH analyses; CAREHF results were inspected before the configuration was formally finalized but did not alter it or any other analytical decision. CAREHF was therefore treated as an external transportability benchmark rather than an untouched confirmatory cohort.

The untouched external confirmation dataset, GSE270788, included 23 paired RNA–ATAC samples(3). Of 46,384 nuclei passing source-level processing, 31,477 were assigned to 9 compatible frozen CardioChrom routes. The remaining 14,907 nuclei were retained as secondary or unmapped observations and were not forced into incompatible routes. Ventricular cardiomyocyte, atrial cardiomyocyte, and neuronal routes had no qualifying nuclei under the frozen mapping rules and were not reconstructed.

SCP3342 contained 48,866 RNA-only nuclei from 43 donors, including 24 nonfailing controls and 19 patients with heart failure with preserved ejection fraction(4). This dataset was used only after the reconstruction models and analytical gates had been frozen.

**Ethical Oversight**

The present study was a secondary computational analysis of deidentified human datasets. The original studies reported institutional review board or ethics committee approval and informed consent as applicable(1-4). No new participants were recruited and no new biological specimens were collected for the present analysis. The University of Arizona Institutional Review Board determined that this secondary analysis did not constitute human subjects research (STUDY00008718); therefore, IRB approval was not required.

**Feature Harmonization and Preprocessing**

RNA profiles were restricted to a fixed universe of 36,100 genes. For each nucleus, RNA counts $x_g$ were library-size normalized and log transformed as

$$X_g = \log\left[1 + \frac{10000x_g}{\sum_i x_i}\right]$$

Chromatin accessibility was represented in a fixed source-derived universe of 285,873 canonical peaks. RNA and ATAC feature spaces were reduced independently to 50 dimensions using truncated singular-value decomposition. Within every training fold, singular-value decomposition was fitted using training donors only and after exclusion of the cell type designated for leave-one-cell-type-out evaluation. Validation, test, and external observations were projected using the corresponding frozen transformations.

Histone-modification signals were mapped to 191,678 nonoverlapping 15-kb genomic bins and normalized using the same library-size log-transformation principle. Separate rank-50 target transformations were fitted for H3K27ac and H3K27me3 using only their respective same-nucleus training observations. All genomic operations crossing multiple bins were chromosome aware and did not extend across chromosome boundaries.

**CardioChrom Latent-State Reconstruction**

CardioChrom was adapted from the latent-space nearest-neighbor strategy developed in the NeurIPS multimodal single-cell integration challenge(5). Twelve leave-one-cell-type-out models were trained. In each fold, all nuclei belonging to one cardiac cell type were excluded from model training, and the corresponding RNA and target-modality latent spaces were fitted from the remaining training donors and cell types.

For each query nucleus, the normalized RNA profile was projected into the frozen 50-dimensional RNA latent space. The 25 nearest training nuclei were identified using Euclidean distance. The predicted target latent state was the uniformly weighted mean of the target-modality latent vectors of these neighbors. Continuous feature-level predictions were obtained by inverse transformation into the frozen ATAC or histone feature space.

No donor identifier, disease label, sex, external-cohort label, or target-modality measurement from the query nucleus was supplied to the predictor. The number of neighbors, distance metric, feature universes, latent dimensions, and reconstruction rules were frozen before external evaluation. Predictions from different cross-validation folds were not averaged for external cohorts.

**Negative and Baseline Comparators**

Three comparator classes were used. First, a within-cell-type RNA-shuffled control disrupted the correspondence between RNA and target-modality states while preserving cell-type composition and the marginal distributions of the input and target features. This comparator assessed whether reconstruction depended on cell-specific RNA information rather than cell-type-average structure.

Second, ATAC predictions were compared with a training-set peak-frequency prior. For each peak, the prior probability was calculated only from eligible training nuclei as

$$p_j = \frac{n_{\text{open},j}+0.5}{N_{\text{train}}+1}$$

Where $n_{\text{open},j}$ was the number of accessible training nuclei at peak $j$ and $N_{\text{train}}$ was the number of eligible training nuclei. This Jeffreys-smoothed prior was frozen before evaluation.

Third, histone reconstruction was compared with a training-set mean profile and with DualHistFormer. DualHistFormer was a shared-encoder Transformer comparator implemented in PyTorch version 2.7.1(6, 7). “Shared encoder” indicates that one RNA encoder generated a common latent representation for the two mark-specific output heads; it does not imply that H3K27ac and H3K27me3 were measured in the same nuclei or pseudo-paired across experiments.

DualHistFormer represented the 50 RNA singular-value components as scalar tokens processed by a 2-layer Transformer with a model dimension of 128, 4 attention heads, a feed-forward dimension of 384, and a 192-dimensional shared state. The H3K27ac head predicted a mark-specific rank-50 latent representation and included an auxiliary loss over 512 sampled 15-kb bins with weight 0.20. The H3K27me3 head predicted its own rank-50 representation and included an auxiliary loss over 512 sampled 225-kb broad-domain blocks with weight 0.25. Latent reconstruction used the smooth L1 loss. Models were trained for 10 epochs using the AdamW optimizer with a batch size of 512, learning rate of $3\times10^{-4}$, weight decay of $1\times10^{-4}$, and gradient-norm clipping at 1.0. Automatic mixed-precision training was enabled on CUDA. For each fold, the best checkpoint was selected using the equal-weight mean of the two mark-specific validation losses and was frozen before testing.

**Internal Evaluation of Chromatin Accessibility Reconstruction**

Internal evaluation used the six held-out FNIH test donors and the cell type excluded from each training fold. Evaluation pairs were fixed before model comparison and were shared by CardioChrom and all comparators. Area under the precision–recall curve was the primary feature-level discrimination metric because chromatin accessibility was sparse; area under the receiver-operating-characteristic curve was secondary. Performance was additionally stratified by training-set peak-frequency quintile to determine whether improvements were restricted to common peaks.

Metrics were summarized at donor and cell-type levels. Donors, rather than nuclei or cell–peak pairs, were treated as the biological replicates. Macro-averaged estimates were calculated so that large cell populations did not dominate the overall result.

**External ATAC Evaluation**

For CAREHF, each of the 12 route-level evaluations used 1,024 cell draws, with 256 naturally sampled peaks per draw, yielding 262,144 cell–peak pairs per route. Across routes, 11,512 unique cells contributed to the benchmark. For rare routes, all available cells were used before additional draws were sampled with replacement. CardioChrom and the comparators were evaluated on identical pairs.

For GSE270788, route assignments were determined using frozen source-to-CardioChrom cell-type mappings. No alternative mapping was introduced after inspection of prediction performance. Sample-specific callable-peak masks and evaluation pairs were fixed before the measured ATAC truth was accessed. The final benchmark comprised 9×1,024×256=2,359,296 cell–peak pairs. Performance was summarized first within each sample and route and then across samples; individual nuclei and peak pairs were not treated as independent biological replicates.

**Histone-State Reconstruction and Evaluation**

H3K27ac and H3K27me3 were reconstructed independently using the same 25-nearest-neighbor latent-state procedure. H3K27ac was evaluated at 15-kb resolution. Because H3K27me3 forms broad domains, its primary global metric was a prespecified domain score integrating chromosome-aware 75- and 225-kb representations.

“Direct reconstruction” denotes evaluation in held-out nuclei for which the corresponding histone mark had been directly measured in the same nucleus. “Global-profile fidelity” denotes agreement between predicted and measured donor-by-cell-type pseudobulk profiles across the complete prespecified genomic-bin universe. It therefore measures broad genomic organization and should not be interpreted as cell-level prediction accuracy or recovery of disease effects.

Six prespecified metrics were used to compare CardioChrom and DualHistFormer. For H3K27ac, these were donor-pseudobulk Pearson correlation across 15-kb bins, area under the precision–recall curve for high-signal 15-kb regions, and Pearson agreement of heart-failure-associated effects among the top 5% of disease-responsive regions. For H3K27me3, these were the 75-/225-kb broad-domain score, area under the precision–recall curve for high-signal 225-kb regions, and disease-effect agreement among the top 5% of disease-responsive regions. The training-set mean and RNA-shuffled predictions were evaluated using the same region sets and aggregation rules.

Spatial organization around cardiac candidate cis-regulatory elements was evaluated using metaprofiles spanning ±150 kb around source-defined active elements for H3K27ac and repressive elements for H3K27me3. Measured, CardioChrom-predicted, and RNA-shuffled profiles were processed identically.

**Disease-Associated Histone Remodeling**

Cross-assay disease-effect analyses were performed within each held-out cell type using the frozen FNIH test donors. The primary contrast combined ischemic and nonischemic heart-failure donors and compared them with nonfailing controls. Ischemic-versus-control and nonischemic-versus-control contrasts were examined as sensitivity analyses.

For each region, the disease effect was the difference between the mean heart-failure and control donor-pseudobulk values. H3K27ac effects were calculated in 15-kb bins. H3K27me3 and measured ATAC were aggregated into chromosome-aware 225-kb blocks.

Directional concordance was defined as the proportion of regions in which the predicted histone effect had the biologically expected direction relative to measured ATAC among regions in the top 5% of absolute measured-ATAC disease effects. H3K27ac and ATAC were expected to change in the same direction. Because H3K27me3 is a repressive mark, its effects were sign reversed before comparison, so a reduction in H3K27me3 corresponded to increased regulatory activity. Effect-size agreement was quantified using Spearman correlation across all regions with finite effects after the same orientation adjustment.

**Regulatory Network and Pathway Analyses**

SCENIC+ version 1.0a2 was applied separately using measured ATAC, CardioChrom-predicted ATAC, and RNA-shuffled ATAC together with the same RNA profiles(8). Nuclei, genes, regulatory regions, transcription-factor universes, and evaluation rules were otherwise held constant. Networks derived from measured ATAC served as the reference. Recovery was evaluated using equal-sized top-ranked TF-to-region and region-to-gene sets so that comparisons were not driven by network size.

For GSE270788, sample aggregation and top-set parameters were selected by evaluating cross-sample recovery within that series. Accordingly, these analyses were interpreted as optimized external transportability analyses and not as fully untouched confirmatory tests.

Positive cardiac cCRE-to-gene links from the source-derived activity-by-contact table were used as the regulatory scaffold(9). Activity-by-contact weights were normalized within each target gene. Gene-set analyses used Gene Set Enrichment Analysis, the 50 Hallmark gene sets, and

2,100 Reactome pathways(10-12). For Supplemental Figure 2A, measured-H3-supported pathways were defined as the top 10 Hallmark and top 50 Reactome pathways ranked by absolute measured-H3 NES. Representative pathways were retained when CardioChrom recovered the measured-H3 direction while the RNA-shuffled control did not and the CardioChrom NES was closer to measured H3. Highly overlapping pathways were collapsed, yielding six ventricular-cardiomyocyte, four endothelial, and three pericyte pathways for visualization. For CAREHF, pathway-level regulatory-state refinement was evaluated by comparing measured ATAC alone with measured ATAC plus virtual H3 against the frozen FNIH measured-H3 pathway reference. Improvement was summarized as the change in pathway NES Spearman correlation, the reduction in median absolute NES error, the change in direction concordance, and the change in measured-supported pathway recall.

**Frozen Deployment to SCP3342**

The frozen CardioChrom-ATAC, CardioChrom-H3K27ac, and CardioChrom-H3K27me3 models were applied to SCP3342 without model refitting, feature reselection, recalibration, or exclusion based on prediction results.

Of 48,866 nuclei, 48,433 were assigned to 10 prespecified routes. These comprised 14,644 cardiomyocytes assigned to the ventricular cardiomyocyte route, 291 adipocytes, 12,562 fibroblasts, 9,216 endothelial nuclei obtained by combining Endothelial1 and Endothelial2, 1,666 endocardial cells, 5,100 pericytes, 3,095 myeloid nuclei obtained by combining Macrophage, Proliferating_macrophage, and Mast_cell, 558 vascular smooth muscle cells, 907 lymphocytes, and 394 neuronal cells. The remaining 433 lymphatic endothelial nuclei lacked an unambiguous frozen route and were left unmapped rather than reassigned. This yielded 30 route-by-modality deployments representing 10 mapped cell types and 3 reconstructed modalities.

All 43 donors were represented in the endothelial, fibroblast, ventricular cardiomyocyte, pericyte, and myeloid compartments. Other routes contained fewer donors according to cell availability. The adipocyte route contained only 291 nuclei, including 16 nuclei from 7 HFpEF donors, and was retained with this limitation explicitly recognized.

**Construction of Donor-Level Regulatory Layers**

Six donor-level analysis layers were constructed within each mapped cell type: measured RNA, CardioChrom-ATAC, CardioChrom-H3K27ac, orientation-adjusted CardioChrom-H3K27me3, a histone composite, and a regulatory composite.

For measured RNA, raw counts were summed across all eligible nuclei from each donor and cell type and normalized as $\log\left[1+\frac{10000x_g}{\sum_i x_i}\right]$ Predicted ATAC profiles were averaged across nuclei from the same donor and cell type.

For reconstructed histone profiles, decoded log-scale values were restricted to the prespecified numerical range of 0 to 20, transformed using expm1, renormalized to 10,000 total units per nucleus, aggregated across nuclei from the same donor, and log transformed. H3K27me3 values were first aggregated into chromosome-aware 225-kb blocks.

Positive source-derived activity-by-contact links were used to aggregate chromatin features to genes. For gene $g$,

$$S_{dg} = \frac{\sum_j w_{jg} X_{dj}}{\sum_j w_{jg}}$$

where $X_{dj}$ was the donor-level value of regulatory feature $j$, and $w_{jg}$ was its positive activity-by-contact weight for gene $g$. Only genes with evaluable, nonzero-variance values in all four primitive layers—RNA, ATAC, H3K27ac, and H3K27me3—were retained for cross-layer analyses.

Each primitive donor-by-gene matrix was standardized gene-wise across donors without using disease labels. H3K27me3 was multiplied by −1 before standardization to express it as an activation-oriented score. The histone composite was the mean of standardized H3K27ac and activation-oriented H3K27me3, followed by gene-wise restandardization across donors. The regulatory composite was the mean of standardized ATAC, H3K27ac, and activation-oriented H3K27me3, followed by the same restandardization.

**Donor-Level Pathway Analysis in SCP3342**

Hallmark and Reactome gene sets were intersected with the evaluable activity-by-contact target-gene universe. Pathways containing 15 to 500 evaluable genes were retained. For each donor, pathway activity was calculated as the mean standardized gene score across pathway members.

The unadjusted disease effect was the HFpEF-minus-control difference in mean donor-level pathway score. Two-sided Welch t tests were used for inference. Effect size was reported as Hedges' $g$,

$$g = J\frac{\bar{X}_{\mathrm{HFpEF}} - \bar{X}_{\mathrm{control}}}{s_p}, J = 1 - \frac{3}{4df-1},$$

where $s_p$ was the pooled standard deviation and $df = n_{\mathrm{HFpEF}} + n_{\mathrm{control}} - 2$.

Sex-adjusted estimates were obtained using ordinary least-squares models containing an intercept, HFpEF status, and a binary sex indicator. HC3 heteroscedasticity-consistent covariance estimates were used. Adjusted analyses required at least 8 evaluable donors with both disease groups and both sex categories represented.

Benjamini–Hochberg correction was applied within each prespecified cell-type-by-layer-by-library family and separately across the complete primary testing universe of 28,248 tests(13). Sex-adjusted *P* values were corrected across the same global universe. A pathway association was classified as robust only when it had a global false discovery rate below 0.05 in the primary analysis, retained a global false discovery rate below 0.05 after sex adjustment, and preserved the full-analysis effect direction in at least 90% of leave-one-donor-out iterations. Family-level false discovery rates were reported as complementary descriptive results. For Figure 5D, endothelial Hallmark pathways with at least one robust layer were ranked by the number of robust layers, followed by the maximum absolute Hedges' $g$ and global false discovery rate, and the 10 highest-ranked pathways were displayed.

**Biosample-Pool Sensitivity Analysis**

All 8 SCP3342 biosample pools contained both HFpEF and control donors. To reduce sensitivity to global pool shifts, a competitive pathway score was calculated for each donor as the mean score of pathway-member genes minus the mean score of the complementary common activity-by-contact target-gene universe. These scores were analyzed using HC3 ordinary least-squares models containing disease status, sex, and biosample pool as fixed effects. Pool-adjusted analyses were treated as post-discovery sensitivity analyses and did not replace or modify the frozen primary robustness gate.

**Secondary Rank-Based Gene-Set Enrichment Analysis**

Secondary enrichment analysis ranked genes using their raw HFpEF-minus-control donor-level disease differences converted to signed percentiles within each cell type and layer. Weighted Gene Set Enrichment Analysis used an exponent of 1, gene-set sizes of 15 to 500, and random seed 20260824(10). These analyses were reported separately from the primary donor-level pathway-score inference.

For descriptive cross-layer state classification, each activity-by-contact-linked cCRE–gene relationship was defined as activated when its reconstructed ATAC and H3K27ac effects were positive and its H3K27me3 effect was negative. A relationship was defined as repressed when ATAC and H3K27ac effects were negative and the H3K27me3 effect was positive. All remaining configurations were classified as mixed. These link-level states were summarized by target gene for visualization.

**RNA-Only Transcription-Factor Activity Analysis**

RNA-only transcription-factor activity was analyzed independently in the SCP3342 **endothelial and fibroblast compartments** using pySCENIC version 0.12.1 (14). The endothelial compartment combined Endothelial1 and Endothelial2 nuclei (9,216 nuclei), and the fibroblast compartment contained 12,562 nuclei. Each compartment included 43 donors: 24 controls and 19 patients with HFpEF.

Genes detected in at least 3 nuclei were retained, and the input transcription-factor universe contained 1,892 factors. GRNBoost2 was used to infer transcription-factor-to-gene relationships, followed by cisTarget motif enrichment and pruning and AUCell regulon scoring(14, 15). Only positive regulons were analyzed. The workflow used random seed 666, a cisTarget rank threshold of 5,000, enrichment AUC threshold of 0.05, normalized enrichment score threshold of 3.0, motif-similarity false discovery rate of 0.001, module thresholds of 0.75 and 0.90, up to 50 top targets, top-regulator settings of 5, 10, and 50, a minimum regulon size of 20 genes, and no dropout mask. AUCell used an AUC threshold of 0.05 and retained the original cell identifiers.

Cell-level AUCell scores were averaged within each donor. For each regulon, donor-level activity was modeled by ordinary least squares with disease status and sex as predictors. The standardized disease effect was defined as

$$\beta_{\text{standardized}} = \frac{\beta_{\text{HFpEF,sex-adjusted}}}{\text{SD}(\text{donor} - \text{level AUCell})}$$

Transcription factors were ranked within each cell compartment by the absolute standardized disease effect. Two-sided t test *P* values and Benjamini–Hochberg false discovery rates were retained as descriptive statistics but were not used as transcription-factor selection gates.

**CardioChrom Transcription-Factor Prioritization**

CardioChrom transcription-factor prioritization was performed without model retraining. Eligible transcription factors were required to have direct transcription-factor-to-gene relationships in the source Table S20 (is_extended = false), corresponding positive motif evidence in Table S14, detectable RNA expression in SCP3342, finite regulatory disease scores for their targets, and at least 5 evaluable direct target genes. Prior heart-failure regulatory networks from Table S19 were not used as a discovery filter or ranking criterion.

For transcription factor $t$, CardioChrom importance was calculated as

$$I_t = \frac{\sum_i w_{ti} \, |R_i|}{\sum_i w_{ti}}$$

where $w_{ti}$ was the Table S20 TF2G_importance_x_abs_rho weight for target $i$, and $R_i$ was the SCP3342 regulatory-composite disease score for that target.

For network visualization, 5 transcription factors with complete motif-supported transcription-factor-to-cCRE-to-gene relationships were displayed for each compartment. Direct Table S20 relationships were linked to the source-derived cardiac activity-by-contact cCRE-to-gene scaffold and SCP3342 activity-by-contact-linked disease effects. cCRE sequences were evaluated using JASPAR 2026 motifs within the canonical 285,873-peak universe at a false-positive rate of $1\times10^{-4}$(16). All supported edges were retained analytically; gene labels were limited only for display clarity. The resulting networks were interpreted as inferred regulatory networks and not as causal networks.

For bidirectional comparison, RNA-only pySCENIC and CardioChrom ranks were converted independently to within-method percentiles. A transcription factor absent from the eligible universe of the comparison method was labeled "not represented" rather than assigned a rank or score of zero.

**Statistical Analysis**

The donor or external sample was the independent biological unit. Nuclei, genomic bins, peaks, genes, pathways, and network edges were not treated as independent biological replicates. Reconstruction metrics were first aggregated within donor or sample and cell type before comparisons across biological units.

All hypothesis tests were two sided unless otherwise specified. Paired external sample-level comparisons used two-sided Wilcoxon signed-rank tests. Disease-associated gene and pathway differences were summarized using Hedges' $g$. Multiple testing was controlled using the Benjamini–Hochberg procedure(13). No nucleus, donor, route, transcription factor, gene, pathway, or prediction was excluded after inspection of its disease association or prediction performance unless it failed a prespecified eligibility criterion. Analyses were performed in Python and R. CardioChrom source code and archived software are publicly available as described below (17).

**Software and Computational Environment**

CardioChrom analyses were performed using Python version 3.11.16 with NumPy version 2.4.6, pandas version 2.3.3, SciPy version 1.17.1, scikit-learn version 1.9.0, and AnnData version 0.10.9. DualHistFormer was implemented in PyTorch version 2.7.1 under Python version 3.11.4. Regulatory-network analyses used SCENIC+ version 1.0a2 under Python version 3.11.8 and pySCENIC version 0.12.1 under Python version 3.10.20. R-based analyses and visualization were performed using R version 4.5.2 and ggplot2 version 4.0.3.

**Data and Code Availability**

Source datasets are available through the access routes reported by the original studies (1-4). CardioChrom source code is available at GitHub under the MIT License. CardioChrom version 1.0.0, the frozen model bundle, and associated release materials are archived at Zenodo (17).

**Supplemental Figure 1. Comparison of CardioChrom and Transformer-based histone reconstruction.**

CardioChrom and the Transformer comparator were evaluated across six prespecified histone-reconstruction metrics. For H3K27ac, metrics included donor-pseudobulk Pearson correlation, high-signal-region AUPRC, and HF-effect Pearson correlation. For H3K27me3, metrics included the broad-domain score, high-domain AUPRC, and HF-effect Pearson correlation. Lines connect CardioChrom and Transformer results for the same metric.

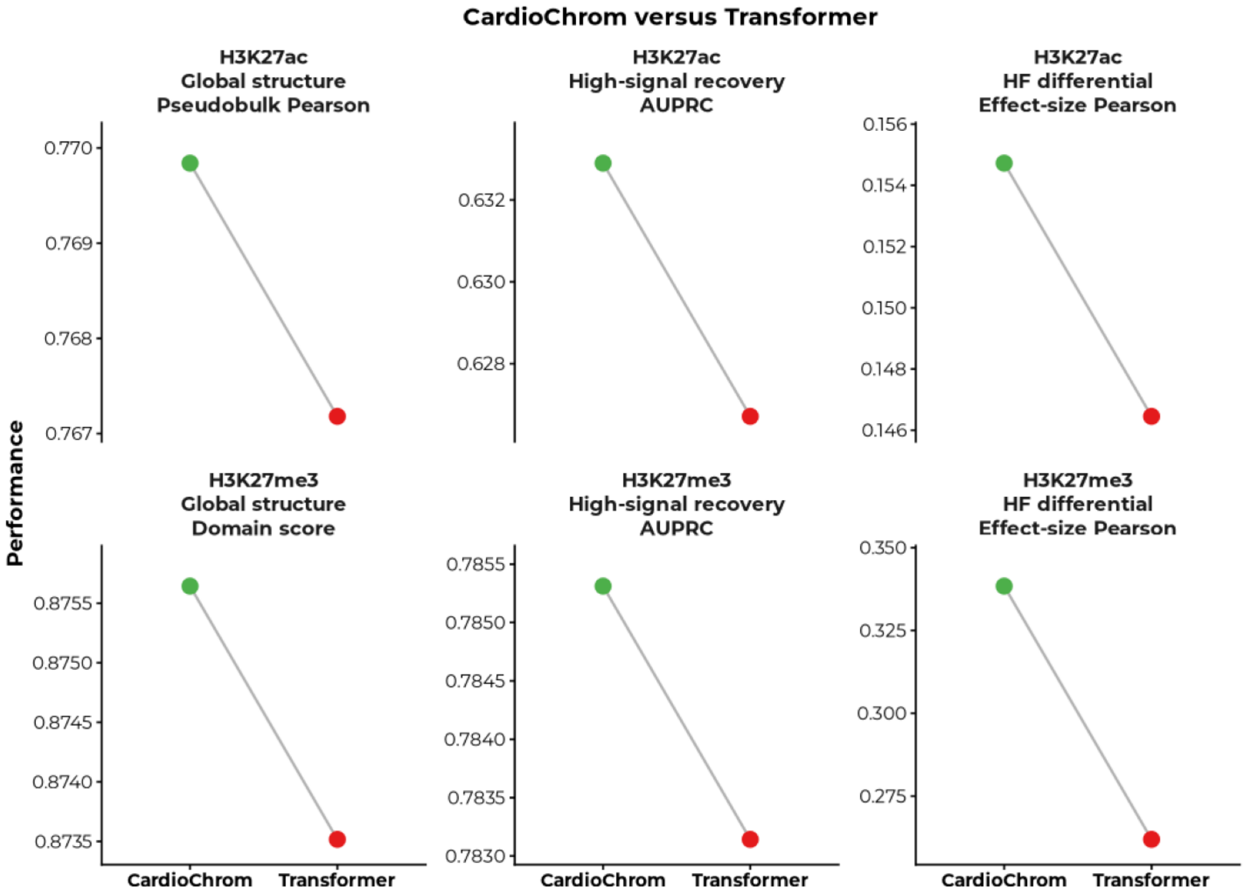

**Supplemental Figure 2. Pathway-level validation of CardioChrom-reconstructed histone states.**

**A,** Recovery of measured-H3-associated disease programs in held-out FNIH ventricular cardiomyocytes (vCM), endothelial cells, and pericytes. Tiles show pathway NES for measured H3, virtual H3, and the RNA-shuffled control.

**B,** Regulatory-state refinement in CAREHF after adding virtual H3 to measured ATAC, evaluated against the FNIH measured-H3 pathway reference in vCM and atrial cardiomyocytes (aCM). Metrics show changes in pathway-pattern similarity, NES error, direction agreement, and pathway recovery for Hallmark and Reactome gene sets. Positive values indicate improvement.

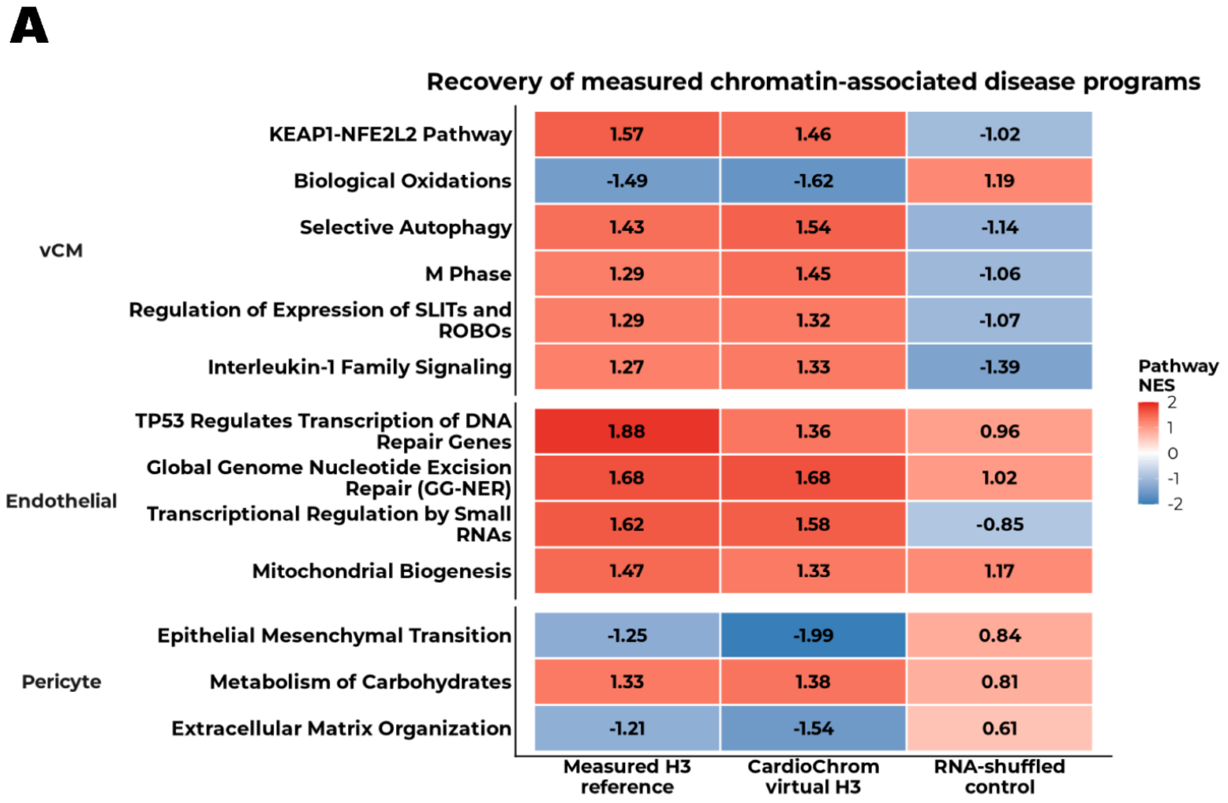
A
Recovery of measured chromatin-associated disease programs
vCM
KEAP1-NFE2L2 Pathway 1.57 1.46 -1.02
Biological Oxidations -1.49 -1.62 1.19
Selective Autophagy 1.43 1.54 -1.14
M Phase 1.29 1.45 -1.06
Regulation of Expression of SLITs and ROBOs 1.29 1.32 -1.07
Interleukin-1 Family Signaling 1.27 1.33 -1.39
Endothelial
TP53 Regulates Transcription of DNA Repair Genes 1.88 1.36 0.96
Global Genome Nucleotide Excision Repair (GG-NER) 1.68 1.68 1.02
Transcriptional Regulation by Small RNAs 1.62 1.58 -0.85
Mitochondrial Biogenesis 1.47 1.33 1.17
Pericyte
Epithelial Mesenchymal Transition -1.25 -1.99 0.84
Metabolism of Carbohydrates 1.33 1.38 0.81
Extracellular Matrix Organization -1.21 -1.54 0.61
Measured H3 reference
CardioChrom virtual H3
RNA-shuffled control
Pathway NES
2
1
0
-1
-2

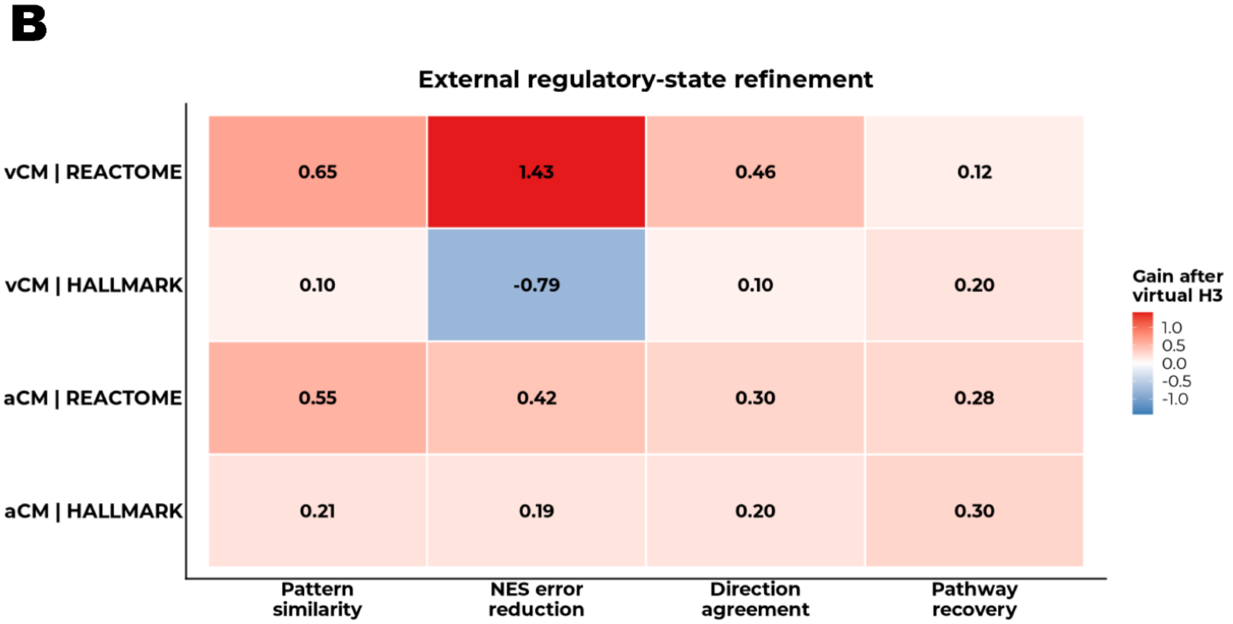
B
External regulatory-state refinement
vCM | REACTOME 0.65 1.43 0.46 0.12
vCM | HALLMARK 0.10 -0.79 0.10 0.20
aCM | REACTOME 0.55 0.42 0.30 0.28
aCM | HALLMARK 0.21 0.19 0.20 0.30
Pattern similarity
NES error reduction
Direction agreement
Pathway recovery
Gain after virtual H3
1.0
0.5
0.0
-0.5
-1.0
